# scTrace+: enhance the cell fate inference by integrating the lineage-tracing and multi-faceted transcriptomic similarity information

**DOI:** 10.1101/2024.11.12.623316

**Authors:** Wenbo Guo, Zeyu Chen, Xinqi Li, Jingmin Huang, Qifan Hu, Jin Gu

## Abstract

Deciphering the cell state dynamics is crucial for understanding biological processes. Single-cell lineage tracing technologies provide an effective way to track the single-cell lineages by heritable DNA barcodes, but the high missing rates of lineage barcodes and the intra-clonal heterogeneity bring great challenges for dissecting the mechanisms of cell fate decision. Here, we systematically evaluate the feature of single-cell lineage tracing data, and then develop an algorithm scTrace+ to enhance the cell dynamic traces by incorporating multi-faceted transcriptomic similarities into lineage relationships via a Kernelized Probabilistic Matrix Factorization model. We assess its feasibility and performance by conducting ablation and benchmarking experiments on multiple real datasets, and show scTrace+ can accurately predict the fates of cells. Further, scTrace+ effectively identifies some important driver genes implicated in cellular fate decision of diverse biological processes, such as the cell differentiation or the tumor drug responses.

## Introduction

Characterizing the dynamics of cell states is crucial for understanding various biological processes, such as tissue development, cell differentiation, disease progression, and drug response^1^. Recent advances in single-cell RNA sequencing (scRNA-seq) technologies enable the cell-level gene expression profiling at a given time point. However, due to the destructive effects on cells caused by omics detection, it is impossible to track the temporal molecular characteristics in any individual cell over time. In response to this limitation, the emergence of single-cell lineage tracing combined with RNA-sequencing (LT-scSeq) offers a complementary approach for investigating cell dynamics, which experimentally introduce unique DNA sequences (lineage barcodes) into cells via viral infection, DNA cassette recombination, or CRISPR-Cas9 genome editing. After the lineage barcoding, the cells are sampled to perform scRNA-seq at different time points. These DNA barcodes are heritable between the ancestral cells and their descendants, so that the clonal relationships or hierarchies can be reconstructed based on single-cell sequencing results^2,3^ (Fig. 1A). These technologies achieve the simultaneous detection of both cell states and cell lineages, and have been widely applied to investigate the cell fate mechanisms under different scenarios, such as hematopoiesis^4^, *C. elegans* embryogenesis^5^, and cancer drug-tolerant persister states^6^. However, some inherent limitations of lineage tracing technologies pose significant challenges. A notable issue is the substantial barcode off-target and missing effects during the experiments of lineage tracing and scRNA-seq, which can result in a considerable proportion of cells not being labeled or not inheriting ancestral barcodes^3,7^. This inefficient labeling may hinder the identification of some critical biological discoveries. Besides, if there is an long time interval between barcode labeling and sequencing, cells sharing the same barcode may have expanded into multiple heterogeneous branches^3,7^. This will lead to diverse cell states within a single clone, as well as confused ancestor-descendant relationships. In parallel to employing lineage tracing experiments for cell lineage reconstruction, some computational approaches are also designed to analyze cell dynamics based solely on transcriptome data^1^, such as pseudo-time analysis^8^ and some optimal-transport based algorithms^9–11^. These approaches prove the values of transcriptomic similarity in inferring cell dynamics when the biological processes satisfy the assumption that cell states change gradually over time (the cells linked across time points have similar gene expression profiles). Hence, integrating the lineage tracing information with transcriptomic similarity holds promise for advancing our understanding of cell dynamics.

**Fig. 1.**
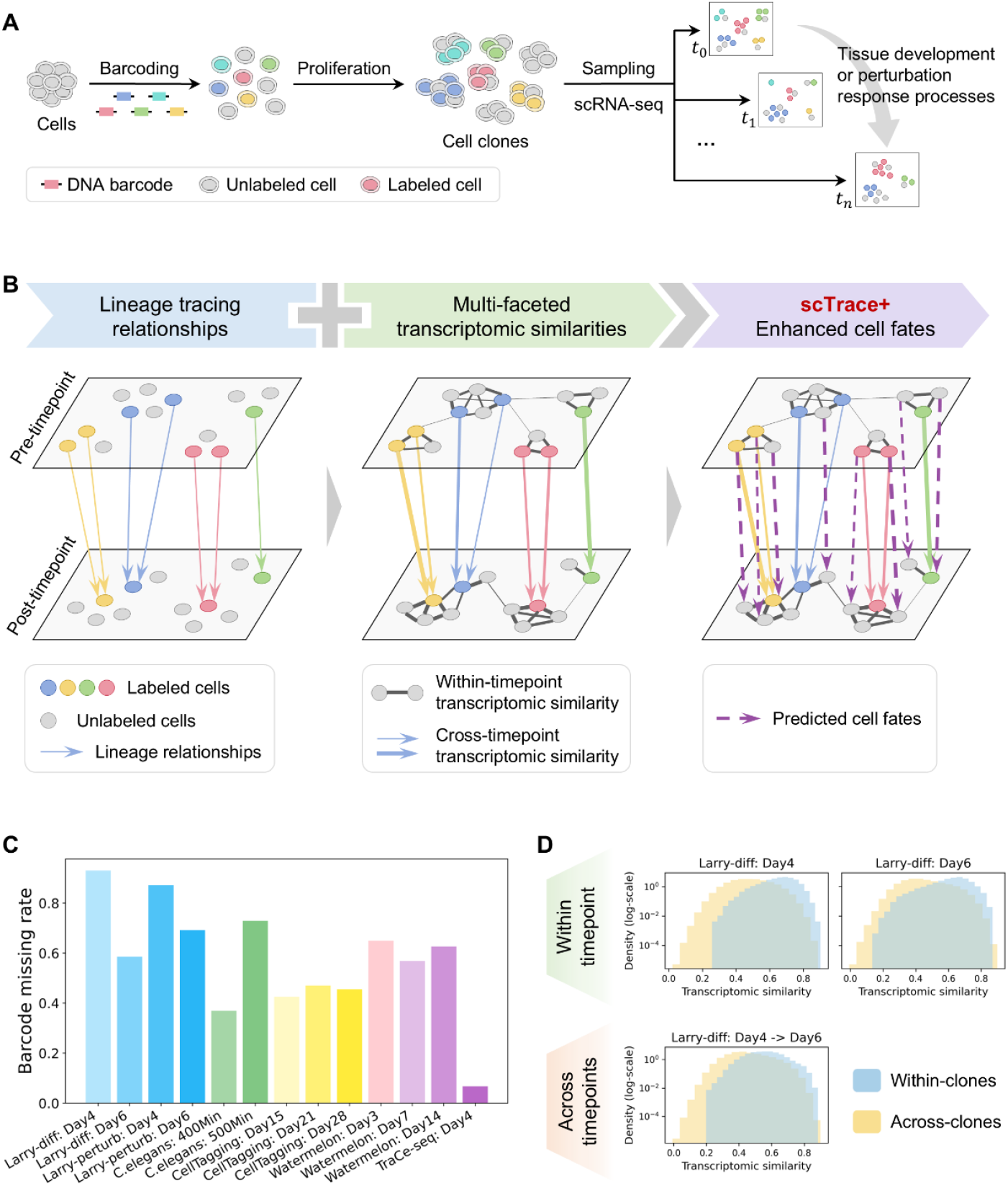
The LT-scSeq experiment and the motivation for scTrace+. **A**. Schematic workflow of time-series LT-scSeq experiments. **B**. The motivation of scTrace+ is to integrate the lineage tracing relationships (left panel) with multi-faceted transcriptomic similarities (middle panel) to overcome the limitations of LT-scSeq data and further enhance the cell fates (right panel). **C**. Barplot showing the high missing rates of inherited lineage barcodes in the cells of multiple LT-scSeq datasets, indicating a significant limitation of LT-scSeq data. **D**. Histograms comparing the transcriptomic similarities among cells within the same clone (blue) versus those across different clones (yellow) in Larry-diff dataset.

Building on this idea, several algorithms have been developed. LineageOT utilizes lineage relationships from posterior time point to adjust cell positions, and then connect the cells across two time points based on optimal transport strategy^12^. Moslin algorithm leverages within-timepoint lineage relationships and across-timepoints expression similarities to reconstruct cellular trajectories through a Fused Gromov-Wasserstein model^13^. Cospar algorithm infers cell dynamics by constraining the coherence and sparsity of transition matrix based on lineage clonal relationships and within-timepoint cell state similarities^7^. While these methods facilitated some valuable discoveries, they do not fully leverage all four types of available information—namely, both lineage tracing relationships and transcriptomic similarities within and across time points (Table S1). These four types of information can provide complementary insights for comprehensively characterizing cell dynamic traces. In detail, the lineage tracing information across time points can be regarded as some fundamental time-series links between cells, while the transcriptomic similarity across time points can infer some gradual transitions in cell states. Within each time point, the lineage relationships and transcriptomic similarities can be utilized as side information to help infer the dynamics of the cells without lineage labeling, based on the principle that the cells belonging to the same clone or exhibiting similar states typically have similar fates.

Here, we first systematically evaluate the missing rates of lineage barcodes in LT-scSeq datasets, and analyze the transcriptomic similarities both within and across cell clones. Then we fully exploit the aforementioned four types of information and specifically develop scTrace+, a computational method designed to enhance (as indicated by the ‘+’ in its name) the single-cell fate inference through integrating lineage tracing information with multi-faceted transcriptomic similarities (both within and across time points) via a Kernelized Probabilistic Matrix Factorization (KPMF) model^14,15^ (Fig. 1B). Our method first integrates the lineage relationships and transcriptomic similarities across time points to balance the heterogeneous cell fate branches and gradual cell state transition. This integration yields a quantification matrix of cell fate transition probability, advancing beyond mere binary ancestor-descendant relationships. Then, we utilize cell-clone and cell-similarity networks within each time point as side-information, and perform low-rank matrix completion to infer more comprehensive cell fate transition probabilities. We prove the performance of scTrace+ through establishing validation sets, conducting ablation experiments, and performing benchmarking comparisons. Furthermore, using the enhanced cell fate transition matrix, we show that scTrace+ effectively facilitates the identification of genes influencing cell fate decisions across various scenarios, including hematopoietic cell differentiation and tumor cell drug response. These biological signals may be ignored when relying solely on experimental lineage tracing data.

## Results

### Data feature evaluation for LT-scSeq

We collected seven publicly available LT-scSeq datasets (Table S2) to conduct the following evaluation. First, for each post-timepoint within these datasets, we calculated the missing rates of inherited lineage barcodes in the cells (Methods, Fig. 1C), and found that more than half of the cells in most datasets did not inherit lineage barcodes from their progenitor cells, indicating significantly inadequate tracking for cells.

Then, to investigate the relationship between transcriptomic similarities and lineage tracing information, we calculated the Pearson correlation coefficients between the gene expression values of any two cells at adjacent time points, and compared the distribution of these similarities both within a single clone and across distinct clones (Methods). The results indicated that the transcriptomic similarities within clones are higher or at least not lower than those observed across clones, regardless of whether the datasets pertain to tissue development or tumor drug response processes (Fig. 1D and S1-S4). This finding inspires us that integrating transcriptomic similarities within-timepoint and across-timepoints may be able to help overcome the high missing rates of lineage barcoding.

### Overview of scTrace+ algorithm

To achieve the objectives outlined above, we specifically designed scTrace+, an algorithm that flexibly integrates lineage tracing information with transcriptomic similarity, both within and across time points, thereby enhancing the characterization for cell dynamic traces. First, we construct a binary matrix to represent lineage relationships between the cells at two time points. To further quantify the strength of these time-series cell links, we calculate the transcriptomic similarity matrix using Pearson correlation analysis and subsequently multiply it onto the lineage relationship matrix in an element-wise way, resulting in a sparse cell transition matrix ***R***_*N*×*M*_, where *N* and *M* denote the number of cells at these two time points. Next, we construct the cell-cell networks 𝒢_*pre*_ and 𝒢_*post*_ for the pre- and post-timepoint, respectively, by integrating the cellular lineage clone relationships and transcriptomic similarities corresponding to each time point. Drawing inspiration from recommender systems^16^, we aim to infer more comprehensive cell dynamic traces through matrix completion. On the one hand, the low-rank property of cell transition matrix across time points can help recovery the missing values from the sparse observations. On the other hand, the cell-cell networks 𝒢_*pre*_ and 𝒢_*post*_ can be treated as side information, suggesting that the cells connected by edges may share similar dynamic traces. These two strategies are incorporated into a KPMF model^14^ to enhance the information regarding single-cell dynamic traces. We decompose the cell transition matrix ***R***_*N*×*M*_ into two low-dimensional latent matrices ***U***_*N*×*D*_ and ***V***_*M*×*D*_, where *D* ≪ *N, M* represents the dimension of latent factors. The independence between any two rows in ***U*** or ***V*** is determined by whether there is an edge between the two corresponding cells in the side information networks 𝒢_*pre*_ and 𝒢_*post*_. Therefore, we define the probabilistic distributions for these variables as follows. The *d* th (*d* ∈ {1, …, *D*}) latent factor (column) in ***U*** and ***V*** is generated from zero-mean Gaussian processes *GP*(***U***|**0, *K***_*U*_) and *GP*(***V***|**0, *K***_*V*_), respectively, where ***K***_*U*_ and ***K***_*V*_ are covariance matrices obtained by applying the Radial Basis Function (RBF) kernel to the graph-embedding matrices of 𝒢_*pre*_ and 𝒢_*post*_. Then, for each observed element ***R***_*n,m*_, we model it through a Gaussian distribution 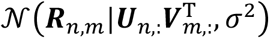, where *σ* represents the standard deviation. Inference for ***U*** and ***V*** is achieved by deriving the log-posterior log *p*(***U, V*** | ***R***, *σ*^2^, ***K***_*U*_, ***K***_*V*_) and seeking the Maximum a Posteriori (MAP) estimation. Finally, we obtain the completed cell transition matrix ***R*\*** by multiplying the matrices ***U*** and ***V***, such that ***R*\*** = ***UV*^T^**. Based on this enhanced cell transition matrix, we can estimate the fate probability for each cell and further identify the driver genes influencing cellular fates (Fig. 2, Methods).

**Fig. 2.**
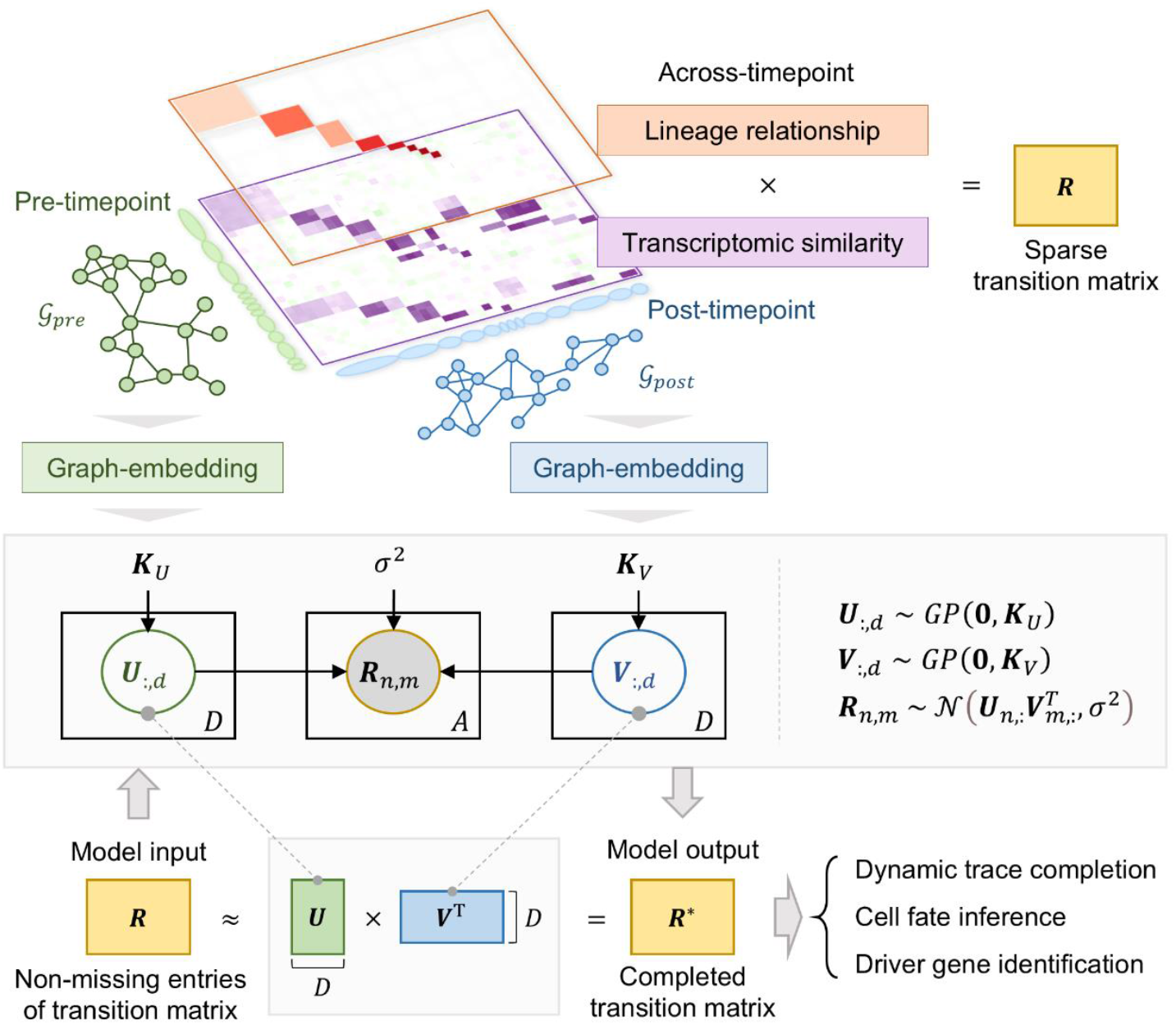
Overview of scTrace+ algorithm. scTrace+ first converts the gene expression matrix and clonal information into a sparse transition matrix ***R*** across time points, as well as cell-cell networks 𝒢_*pre*_ and 𝒢_*post*_ within time points. The networks 𝒢_*pre*_ and 𝒢_*post*_ are then transformed into graph kernels through representation learning. Next, scTrace+ decomposes the sparse matrix ***R*** into low-dimensional matrices ***U*** and ***V***. The completed transition matrix ***R*\*** is derived from the product ***UV***^***T***^ and is utilized for downstream analyses, including dynamic trace completion, cell fate inference, and driver gene identification.

### Performance evaluation of scTrace+

We conducted a comprehensive evaluation of scTrace+ on multiple LT-scSeq datasets (Fig. 3). Firstly, we divided the non-missing entries of the cell transition matrix ***R*** into training and validation sets, and then inferred the completed matrix ***R*\*** via scTrace+ based on training set, with the entries in the validation set masked to facilitate an assessment of predictive performance.

**Fig. 3.**
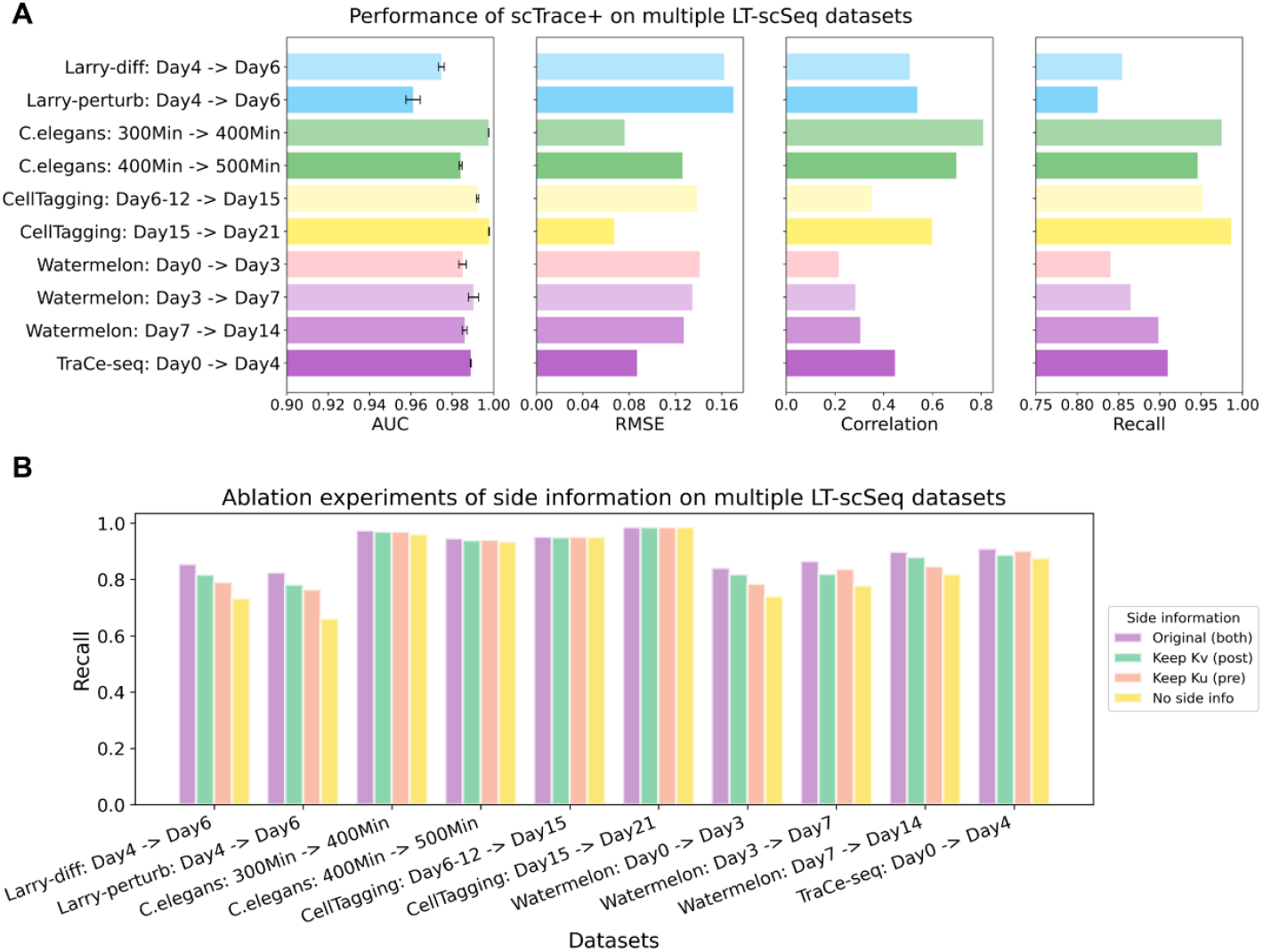
Performance evaluation of scTrace+ on multiple datasets. **A**. Barplots summarizing the prediction performance of scTrace+ on different datasets. The leftmost panel displays the AUC scores on validation sets. The experiment was repeated five times, and the length of the bars along with the error lines represent the mean and standard deviation, respectively. The other three panels showcase the RMSE (Root Mean Square Errors), correlation, and the recall rates of scTrace+ on the validation sets when predicting the non-missing entries, respectively. **B**. Ablation experiments evaluating the influence of side information on multiple datasets. For each dataset, the barplot from left to right represents the recall rate under four scenarios: 1) incorporating all side information, 2) retaining side information 𝒢_*post*_, 3) retaining side information 𝒢_*pre*_, and 4) no side information.

Next, we compared the entries in the completed matrix ***R*\*** with those in the raw binary lineage relationship matrix, and calculated AUC (Area Under the Receiver Operating Characteristic curve) scores to quantify the predictive performance of scTrace+ (Methods). We observed that the AUCs scores were generally larger than 0.95 (the leftmost panel in Fig. 3A), indicating that scTrace+ can always maintain a high recall for the raw lineage relationship entries (high true positive rate), no matter whether the enhancement rates were high or low (represented by varying false “positive” rates due to different classification threshold settings).

By comparing the non-missing entries in the validation set of matrix ***R*** (considered as ground truth) against the predicted values at corresponding positions in the completed matrix ***R*\***, we found that scTrace+ achieved not only low Root Mean Square Errors (RMSE) and high correlation between ground truth and predictions, but also high recall rates for strong positive entries (the right three panels in Fig. 3A, Methods). These results proved that scTrace+ predicted the masked observations (validation set) accurately and enhanced the LT-scSeq data reliably.

Besides, we performed ablation experiments to evaluate the contributions of incorporating side information. We trained the models under four distinct ablation scenarios: 1) integrating side information from both pre- and post-timepoints; 2) using only post-timepoint information; 3) using only pre-timepoint information; and 4) excluding all side information. By testing these models on the validation sets and calculating the recall rates, we found that integrating all of the side information yielded the best performances (Fig. 3B), suggesting that multi-faceted transcriptomic similarity information contributes greatly to the performance of scTrace+.

### scTrace+ facilitates the prediction and dissection of cell fates in hematopoiesis

We applied scTrace+ to two published datasets of mouse hematopoietic progenitor cells with LARRY lentiviral barcode library tagging^4^. The first dataset, referred to as “Larry-diff”, captured cells on day 4 and 6 of the differentiation processes, while the second dataset, termed “Larry-perturb”, sampled cells on day 4 and day 6 after the cytokine perturbation^4^. Both datasets elucidated the differentiation dynamics from hematopoietic progenitor cells into some mature cell types, such as neutrophils and monocytes (Fig. 4A and S5A). Notably, in the Larry-diff dataset, nearly 56.5% of cells were not labeled by the lineage barcodes (Fig. 4B), and only 22% of the cells on day 6 could be traced back to their progenitors based on the lineage barcoding information (Fig. 4C). After the cell fate enhancement by scTrace+, this traceable fraction increased by nearly two-fold (Methods, Fig. 4C). Fig. 4D presents a visualization of the proportions of cell fates that differentiated into neutrophils, monocytes and other cell types before and after the enhancement provided by scTrace+. Similar results were also observed in the Larry-perturb dataset (Fig. S5B and S5C).

**Fig. 4.**
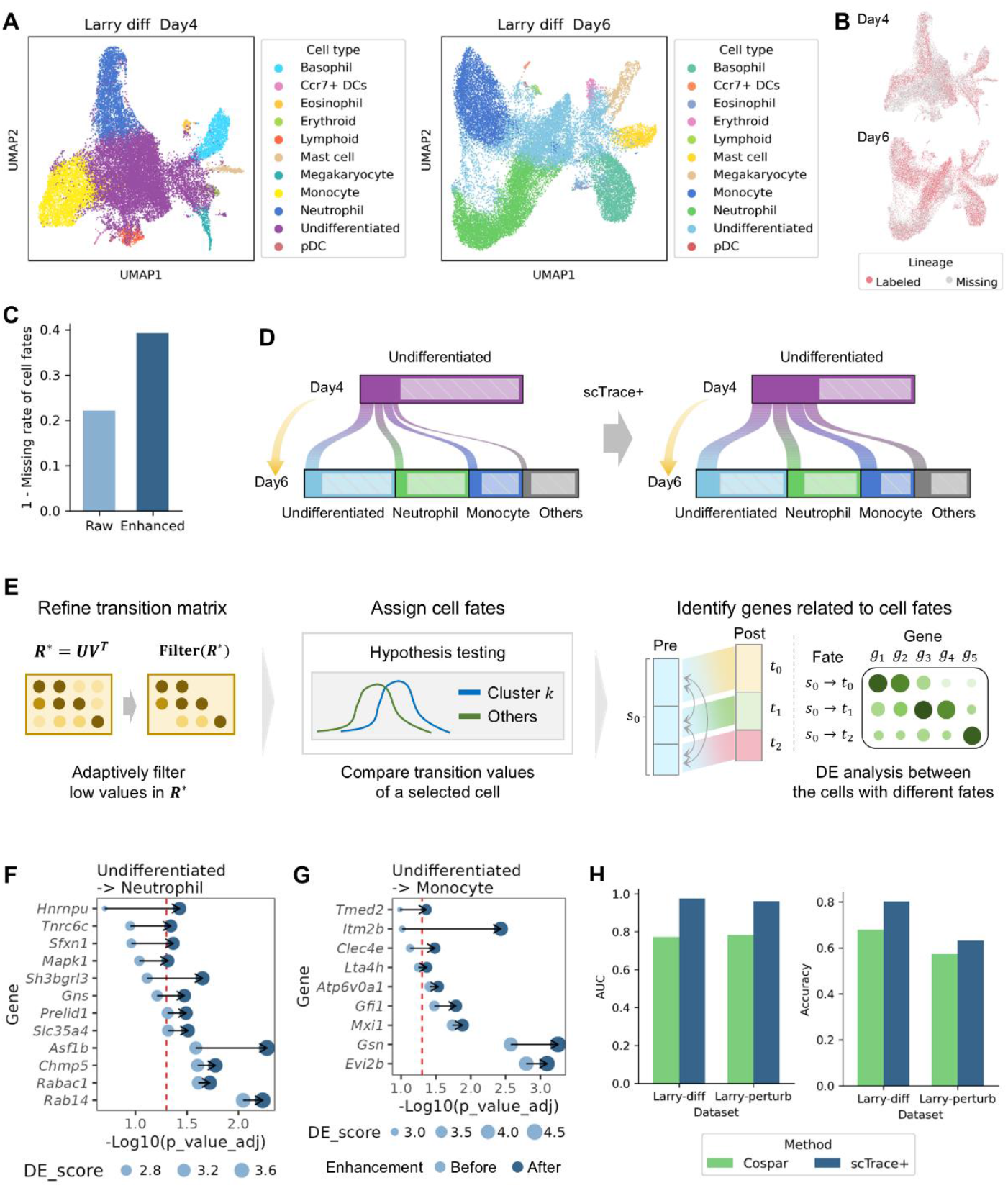
scTrace+ facilitates the prediction and dissection of cell fates in hematopoiesis. **A**. UMAP (Uniform Manifold Approximation and Projection) plots of the cells in Larry-diff dataset at days 4 and 6. The color indicates the cell type label annotated in the original study. **B**. UMAP plots showing the lineage barcode labeling results for the cells in Larry-diff dataset. **C**. The missing rates of cell fates before (light blue) and after (dark blue) the enhancement of scTrace+ in Larry-diff dataset. **D**. Sankey plots showing the dynamic fate relationships between undifferentiated cells and their descendants (neutrophils, monocytes and other cell types) before (the left panel) and after (the right panel) the enhancement by scTrace+. The width of each rectangle corresponds to the number of cells. The shaded areas in light grey represent the cells without fate labeling. **E**. A schematic diagram illustrating the approach for refining cell-cell transition matrix, assigning cluster-level fates to cells, and identifying differentially expressed (DE) genes between the subclusters exhibiting different fates. **F-G**. Bubble plots showing the DE genes with increased statistical significance after the enhancement of cell traces by scTrace+. The DE analyses are conducted between the undifferentiated cells remaining in an undifferentiated state and those transitioning to neutrophils (**F**) or monocytes (**G**). The color of points indicates the types of before (light blue) and after (dark blue) enhancement. The size of points indicates the DE scores calculated by Wilcoxon rank- sum test in Scanpy. The x-axis represents − log_10_(*p*_*value*) of the DE genes. The red dashed line indicates a significance threshold of *p*_*value* **=** 0.05. **H**. Barplots showing the AUC scores and accuracy rates of the fates predicted by Cospar (green) and scTrace+ (dark blue) on the Larry-diff and Larry-perturb datasets.

According to the enhanced cell-cell transition matrix, we assigned cell-type level fate probabilities to each cell and conducted differential expression (DE) analysis to compare cells maintaining undifferentiated states against those differentiating into neutrophils or monocytes (Methods, Fig. 4E). For comparative purposes, we also generated results based solely on the raw lineage information for cell fate assignment and DE gene identification. We found that some genes were newly identified or exhibited increased statistical significances after the enhancement of cell traces (Fig. 4F and 4G). For example, *Mapk1* has been reported as necessary for neutrophil chemotaxis^17^, and *Sh3bgrl3* plays a critical role in regulating various biological processes, such as signal transduction and cell differentiation^18^ (Fig. 4F). Additionally, *Clec4e* and *Lta4h* were also reported to be highly expressed in some monocyte subsets^19,20^ (Fig. 4G). These findings underscore the value of scTrace+ in more comprehensively dissecting the underlying molecular mechanisms governing cell fate decisions.

Furthermore, we assessed the prediction performance of scTrace+ by benchmarking it against Cospar^7^, a recent state-of-the-art method. In our evaluation, we treated the raw lineage information as part of the true time-series relationships among cells, and calculated AUC score to measure the extent to which the entries in the cell-cell fate transition matrix inferred from the raw lineage information were predicted by scTrace+ or Cospar (Methods). Next, we established a set of reasonable cell-type level fates based on the prior biological knowledge and calculated accuracy rates to assess whether the inferred dynamic relationships among cell types belonged to this pre-defined fate set (Methods). The results indicated that scTrace+ achieved superior recall and accuracy rates compared to Cospar (Fig. 4H). Besides, according to the cell-cell transition matrices inferred by raw lineage information, scTrace+ and Cospar, we derived the cell-type level fate proportions for each cell, and found scTrace+ more stably maintained similar fate possibilities to that from raw lineage information (Methods, Fig. S6).

### scTrace+ uncovers the mechanisms underlying heterogeneous drug responses of tumor cells

We applied scTrace+ to a dataset of tumor drug response^21^, which utilized TraCe-seq technology to track heterogeneous clonal responses of lung cancer cell lines (PC9) subject to anticancer drug Erlotinib, an EGFR kinase inhibitor. The cells were labeled by lentivirus-based barcodes and sampled at two time points: day 0 (pre-treatment) and day 4 (post-treatment), for subsequent scRNA-seq. Initially, we conducted clustering analyses on the cells from each timepoint separately (Fig. 5A). Then, we employed scTrace+ to enhance the cell dynamic traces, resulting in an increase in the fraction of cells with inferred fate from 75.3% to 81.3% (Fig. 5B). And the inferred cell fates generally maintained consistent proportions of the cells flowing to different post-clusters with that before enhancement (Fig. 5C).

**Fig. 5.**
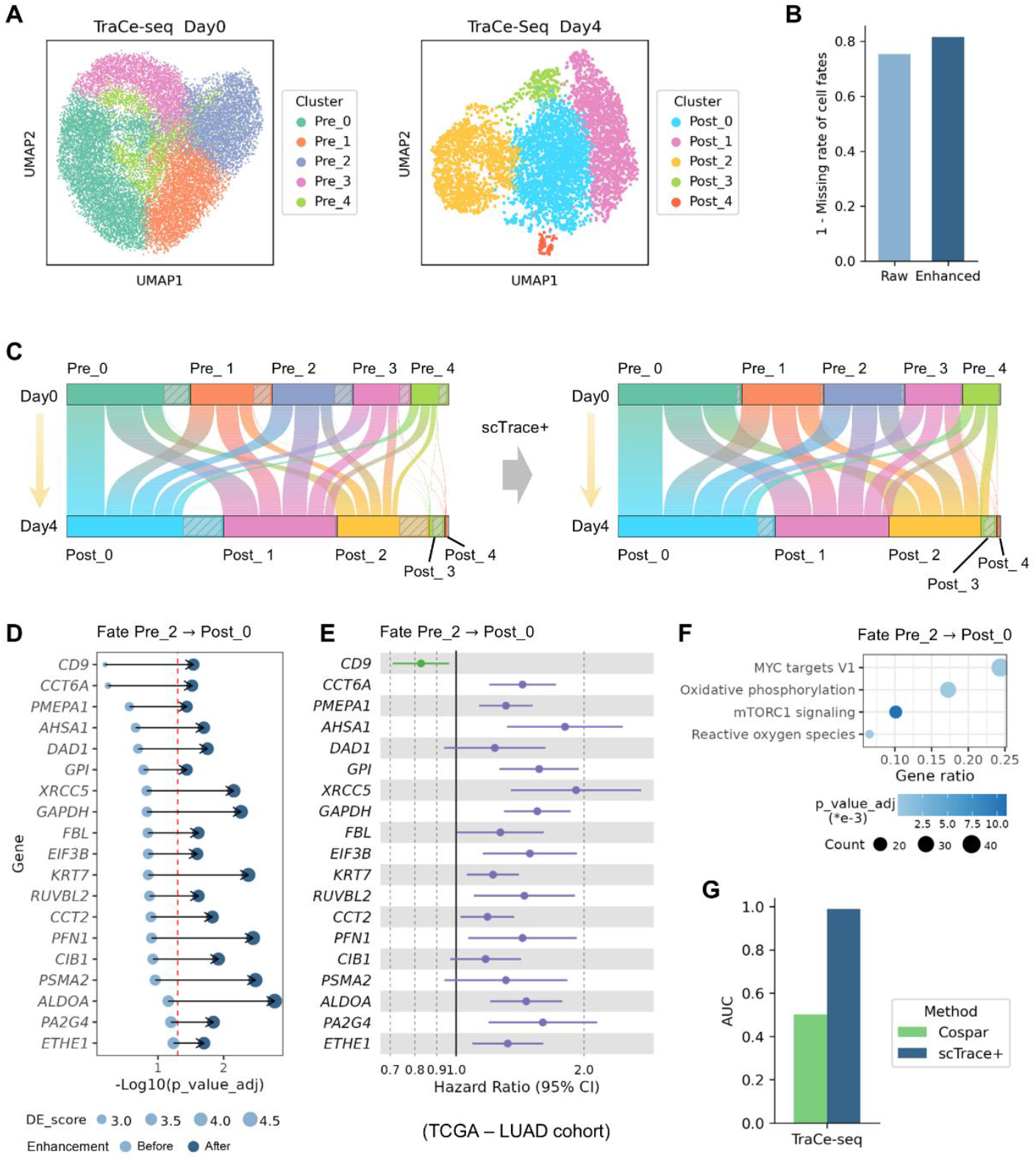
scTrace+ uncovers the mechanisms underlying heterogeneous drug responses of tumor cells. **A**. UMAP plots of the cells in TraCe-seq dataset at days 0 and 4, with color indicating the clustering results. **B**. The missing rates of cell fates before (light blue) and after (dark blue) the enhancement of scTrace+. **C**. Sankey plots showing the dynamic fate relationships between pre-clusters and post-clusters before (the left panel) and after (the right panel) the enhancement by scTrace+. The width of each rectangle corresponds to the number of cells. The shaded areas in light grey represent the cells without fate labeling. **D**. Bubble plots showing the DE genes with increased statistical significances after the enhancement of cell traces by scTrace+. The DE analyses are conducted between the cells in Pre_2 cluster that flowing to Post_0 and other post-clusters. The color of points indicates the types of before (light blue) and after (dark blue) enhancement. The size of points indicates the DE scores calculated by Wilcoxon rank- sum test in Scanpy. The x-axis represents − log_10_(*p*_*value*) of the DE genes. The red dashed line indicates a significance threshold of *p*_*value* **=** 0.05. **E**. A forest plot showing the hazard ratios (HR) of the identified DE genes in TCGA-LUAD (lung adenocarcinoma) cohort. The solid line indicates HR=1. CI, confidence interval. **F**. A bubble plot showing the pathway enrichment analysis results for the DE genes of fate Pre_2→Post_0 after enhancement. The colors of points indicate the adjusted *p*_*value*, and the size of points reflects the number of genes belonging to the corresponding pathway. **G**. Barplots showing the AUC scores of the fates predicted by Cospar (green) and scTrace+ (dark blue) on the TraCe-seq dataset.

Next, we used the methods illustrated in Fig. 4E to assign a cluster-level fate to each cell at day 0 and identify DE genes between the cell subsets exhibiting distinct fates within each pre-treatment cluster. We found the enhancements provided by scTrace+ facilitated the identification of some tumor-associated genes that would have been ignored if solely relying on the raw lineage information. For example, leveraging the enhanced cell fate outcomes led to the identification of some novel DE genes in the transition of cells flowing from cluster Pre_2 to Post_0 (Pre_2→Post_0) (Fig. 5D). Many of these genes have been previously reported in relation to EGFR signal transduction (such as *CD9*^22^), tumor progression (*CCT6A*^23^, *PMEPA1*^24^) and other relevant tumor behaviors. Further, we collected bulk transcriptome and survival data from the TCGA-LUAD (lung adenocarcinoma) cohort, and utilized Cox proportional hazards regression to analyze the impact of these genes on patient survival (Methods). The analysis revealed a significant correlation between most of these genes and poor prognosis (Fig. 5E), highlighting their potential as therapeutic targets. Additionally, these newly identified DE genes enriched on some pathways associated with tumor proliferation and metabolism, including MYC targets, oxidative phosphorylation, and reactive oxygen species. Notably, the enhancements from scTrace+ enabled the identification of the high enrichment on mTORC1 signaling pathway (Fig. 5F and S7), which has been implicated as one of the mechanisms of acquired resistance to EGFR inhibitors in non-small cell lung cancer^25^.

Moreover, we benchmarked scTrace+ against Cospar using this dataset. Our findings indicated that the cell fates predicted by scTrace+ more accurately maintained the raw lineage relationships compared to those derived from Cospar (Methods, Fig. 5G). And the cluster-level fate probabilities inferred by scTrace+ exhibited a closer resemblance to the information obtained from the raw lineage data (Fig. S8).

## Discussion

In this study, we proposed an algorithm called scTrace+ to comprehensively characterize single-cell dynamic traces by integrating both lineage tracing information and multi-faceted transcriptomic similarity within- and across-timepoints. Specifically, the lineage relationships across-timepoints provide some basic time-series links between cells. Integrating the cell-cell transcriptomic similarities across-timepoints can help balance heterogeneous cell fate branches and gradual cell state transition. The cell-cell networks derived from the clonal relationships and transcriptomic similarities within each time point facilitate the inference the dynamics of the cells lacking lineage labels. In scTrace+, these strategies are incorporated into a flexible framework of matrix completion on graphs, utilizing a KPMF model.

The underlying assumption of scTrace+ is that the cells exhibiting similar molecular states or originating a common clonal lineage usually tend to display similar dynamic traces. By collecting multiple LT-scSeq datasets under various scenarios and analyzing their data features, we demonstrated that this assumption generally holds true. Especially for the normal biological processes, such as tissue development and cell differentiation, the transcriptomic similarities within clones are significantly higher than those observed across different clones, reflecting a gradual transition of cell states. While for the tumor-associated processes, such as tumor drug perturbation responses, this phenomenon is a little weak. We reasoned that the inherently high heterogeneities of tumor cells and the pronounced external perturbation may bring some randomness and unpredictability into tumor cell fates. Future exploration may benefit from identifying perturbation-associated driver regulators of cell fates based on observed lineage relationships, thereby enabling more accurate predictions of cell dynamics.

By applying scTrace+ to multiple real datasets, we firstly divided the data into training and validation sets to prove its performance and conducted ablation experiments to assess the effect of incorporating side information. Subsequently, we focused on two datasets involving hematopoietic differentiation and lung cancer cell line drug response processes as case studies to evaluate the utility of scTrace+ in elucidating biological mechanisms. Compared to the analyses using raw lineage relationships, we found the cell dynamic traces enhanced by scTrace+ facilitated the identification of some additional genes and pathways. In the context of hematopoietic differentiation, these findings contribute to a deeper understanding of fate determination and function regulation in different cell types. In the tumor drug response scenario, our findings also revealed some molecular mechanisms underlying heterogeneous drug responses and poor inhibition for tumor cells, which may provide some novel insights for exploring potential therapeutic targets. Furthermore, we compared the performance of scTrace+ with that of a recent state-of-the-art method, Cospar, and demonstrated that the cell fates inferred by scTrace+ exhibited greater consistency with biological prior knowledge, and better maintained the original lineage relationships.

The framework of scTrace+ is highly flexible and scalable. Currently, it integrates all of the lineage and transcriptome information no matter within or across time points. As technological advancements continue, additional information may be incorporated to further enhance the characterization for cell dynamic traces. For example, cumulative barcoding technologies can reconstruct the hierarchical cell lineage tree, so that the distances between cells along the tree can be used as side information to help infer cell dynamics. Single-cell multi-omics technologies offer a more comprehensive characterization of cell states; hence the multi-omics cell-cell similarities can also be integrated to further enhance the lineage tracing information. Moreover, combining the recent spatial transcriptome sequencing with the lineage tracing technologies may enable the directly recording of the spatiotemporal information for cells, so that the spatial neighbor relationships can also be incorporated into the scTrace+ framework to comprehensively dissect the state changes of cells in temporal and spatial dimensions. We believe that the ongoing improvement in lineage tracing technologies and computational algorithms will yield deeper insights into complex biological processes.

## Availability of data and materials

The datasets supporting the conclusions of this article are publicly available. The accession number of the datasets are listed in the Table S2. The Python package implementing scTrace+ and its tutorial material are freely available at https://github.com/czythu/scTrace.

## Acknowledgements

This work was supported by funding from the National Key Research and Development Program of China (Nos. 2020YFA0712403 and 2021YFF1200901), the National Natural Science Foundation of China (NSFC) (Nos. 62133006 and 92268104), the Tsinghua University Initiative Scientific Research Program (No. 20221080076), and the China Postdoctoral Science Foundation (No. 2022M721839).

## Author contributions

J.G. conceived this study. W.G. designed the methods. W.G. and Z.C. performed the analyses and drew the figures. J.G, W.G., Z.C., and X.L. discussed the methods and results. J.H and Q.H. provided some suggestions. Z.C. collected the datasets. W.G., Z.C. and J.G. wrote the manuscript. J.G. supervised the project.

## Methods

### Dataset collection and preprocessing

We apply scTrace+ to seven publicly available LT-scSeq datasets, including developmental datasets and tumor drug response datasets (Table S2). The pre- processing for these LT-scSeq datasets is performed based on the Scanpy^26^, Seurat^27^ or scCancer^28,29^ packages. First, we perform basic quality control steps for scRNA-seq data to filter low-quality cells. Then, we calculate the relative gene expression values by conducting data normalization and log transformation. Next, the union set of the highly variable genes of the data at multiple time points is finally selected for analysis. Furthermore, after performing data centering and scaling, we conduct principal components analysis (PCA) and uniform manifold approximation and projection (UMAP) with parameters n_neighbors = 10 and n_pcs = 40 by default. During the dimension reduction steps, the connectivities of cells are estimated and will be used for further analysis.

### Feature assessment for single-cell lineage-tracing data

#### Missing rates of lineage barcodes

To quantitatively evaluate the sparsity of cross-timepoint lineage relationships in LT-scSeq datasets, we first extract the clonal information from two adjacent time points, comprising *N* and *M* cells, respectively. We then construct a binary adjacency matrix ***L***_*acr*_ ∈ ℝ^*N,M*^, in which the entries indicate whether two cells originated from the same clone. Next, for the post-timepoint, we count the number of cells *m* that inherited lineage barcodes from the cells at pre-timepoint, specifically by determining the number of columns in the binary matrix ***L***_*acr*_ with a column sum greater than 0. Consequently, we define the missing rate of inherited lineage barcodes as 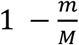 (Fig. 1C).

#### Comparison between transcriptomic similarity and lineage information

Generally, the cellular dynamic processes are characterized by gradual changes in molecular states, with a little randomness introduced by particular event or external perturbation. To evaluate this phenomenon in LT-scSeq data, we calculate the similarity between the gene expression profiles of any two cells, both within- and across-timepoints, using the Pearson product-moment correlation coefficients. Then, we compare the distribution of similarity scores between pairs of cells from the same clone and pairs from different clones, utilizing histograms for visualization (Fig. 1D and S1-S4).

### scTrace+: Kernelized Probabilistic Matrix Factorization for data enhancement

The data enhancement algorithm scTrace+ for LT-scSeq is designed based on a Kernelized Probabilistic Matrix Factorization (KPMF) model^14,15^. Due to the high missing rates of lineage barcodes, the cell transition matrix derived from lineage relationships across time points is highly sparse. Thus, we model the enhancement of LT-scSeq data as the completion of this cell transition matrix. Considering that the possible fates of cells are limited, we think the cell transition matrix is low-rank.

Therefore, we utilize a probabilistic matrix-factorization based strategy to decompose the cell transition matrix into two low-dimensional latent matrices, so that the missing entry can be predicted through the inner product of corresponding latent vectors. Besides, scTrace+ fully integrates the cell-cell networks of transcriptomic similarities and clonal relationships within each time point as side information, so that the fates of cells can be transferred to their neighbor cells and further help the prediction of cell traces. We transform the side information into covariance matrices by utilizing a graph kernel function. Then the covariance matrices are used to constrain the dependency relationships between the latent representations of cells in their prior Gaussian process distributions.

#### Notations

Denoting the number of cells at the pre- and post-timepoint as *N* and *M*, and the number of latent factors as *D*, we define the cell lineage graphs *L* **=** [***L***_*pre*_, ***L***_*post*_, ***L***_*acr*_] and transcriptomic similarity graphs *S* **=** [***S***_*pre*_, ***S***_*post*_, ***S***_*acr*_] as the inputs:

***L***_*pre*_ ∈ ℝ^*N,N*^: a square adjacency matrix. If the two cells at the pre-timepoint originated from the same clone, the corresponding matrix entry is set to 1; otherwise, it is set to 0.

***L***_*post*_ ∈ ℝ^*M,M*^: a square adjacency matrix. If the two cells at the post-timepoint originated from the same clone, the corresponding matrix entry is set to 1; otherwise, it is set to 0.

***L***_*acr*_ ∈ ℝ^*N,M*^: an adjacency matrix cross two time points. If the two cells at adjacent time points originated from the same clone, the corresponding matrix entry is 1; otherwise, it is set to 0.

***S***_*pre*_ ∈ ℝ^*N,N*^ : a transcriptomic similarity matrix of cells at the pre-timepoint, generated by “pp.neighbors” function in the Scanpy package^26^.

***S***_*post*_ ∈ ℝ^*M,M*^: a similarity matrix of cells at the posterior time point, generated by “pp.neighbors” function in the Scanpy package.

***S***_*acr*_ ∈ ℝ^*N,M*^: a similarity matrix of cells across two time points, which is calculated by the Pearson correlation analysis between the gene expression values of any pair of cells across time points.

Next, we derive three variables storing key information for data enhancement from the six input graphs as follows:

***K***_*U*_ ∈ ℝ^*N,N*^ : a side information graph, a kernel matrix modeling the cell-cell relationships at the pre-timepoint, obtained by integrating ***L***_*pre*_ and ***S***_*pre*_ (see the “*Cell covariance matrix generation by kernel transformation*” section for details).

***K***_*V*_ ∈ ℝ^*N,N*^ : a side information graph, a kernel matrix modeling the cell-cell relationship at the post-timepoint, obtained by integrating ***L***_*post*_ and ***S***_*post*_ (see the “*Cell covariance matrix generation by kernel transformation*” section for details).

***R*** ∈ ℝ^*N,M*^: a transition matrix across time points, obtained by element-wise multiplication of ***L***_*acr*_ and ***S***_*acr*_.

Finally, we define the outputs of scTrace+ as follows:

***U*** ∈ ℝ^*N,D*^: a latent matrix at the pre-timepoint, obtained by matrix factorization.

***V*** ∈ ℝ^*M,D*^: a latent matrix at the post-timepoint, obtained by matrix factorization.

***R*\*** ∈ ℝ^*N,M*^: a completed transition matrix, obtained by multiplication of ***U*** and ***V***^***T***^.

Briefly, we approximate the non-missing entries in the sparse transition matrix ***R*** by multiplying the latent matrices ***U*** and ***V***^***T***^; meanwhile, the missing entries are also predicted. In this way, the LT-scSeq data can be enhanced.

#### Cell covariance matrix generation by kernel transformation

To fully integrate the side information, we firstly derive the weighted average of the cell-cell similarity matrix ***S***_***x***_ and clone adjacency matrix ***L***_***x***_ within each time point ***x*** (subscript ***x* =** *pre* and *post*, separately). We set the weight to 0.5 for enhancing developmental datasets with clear fate pattern, while we reduce the weight of transcriptomic similarity to 0.25 for enhancing tumor drug response datasets with higher fate randomness. The constructed cell-cell networks are named as 𝒢_*pre*_ and 𝒢_*post*_.

To reduce the redundant information in high-dimensional side information matrices, we calculate a low-dimensional embedding for each cell in 𝒢_*pre*_ and 𝒢_*post*_. We apply node2vec^30^, a continuous feature learning algorithm to maximize the likelihood of maintaining the neighborhood relationship of cells (the Node2vec package, 32 dimensions by default).

Then, we convert the low-dimensional graph-embedding matrices of 𝒢_*pre*_ and 𝒢_*post*_ into covariance matrices ***K***_*U*_ and ***K***_*V*_ respectively through a kernel function. In scTrace+, we select the Radial Basis Function (RBF) kernel (“rbf_kernel” in the Sklearn package) to quantify the within-timepoint cell-cell similarity. For subsequent calculations, the inverse of the covariance matrix is used.

#### KPMF model construction

As illustrated in Fig. 2, each non-missing entry ***R***_*n,m*_ in the sparse transition matrix ***R*** (*δ*_*n,m*_ **=** 1) is determined by a Gaussian distribution with the mean equaling to the inner product of ***U***_*n*,:_ and ***V***_*m*,:_ and constant variance *σ*^2^:

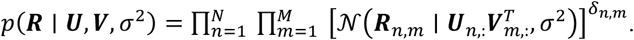

We design the prior distribu^*N*^tion of each column in the latent matrix ***U*** or ***V*** as a zero-mean Gaussian process with a covariance matrix. Notably, to capture correlations among cells within time point, the latent matrices ***U*** and ***V*** are generated in a column-wise manner with covariance matrices ***K***_*U*_ and ***K***_*V*_:

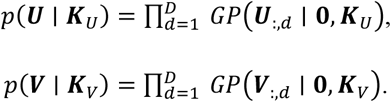

In this way, each factor (column) is independent of all other factors, while the latent representations of cells (rows) are correlated of each other through the covariance matrices. If two cells are linked in the cell-cell network, their corresponding latent factors would be similar after training. Hence, the fate transition probabilities of the cells without lineage labeling can be predicted according to the lineage dynamics of other cells they linked.

#### Deriving log-posterior and minimizing loss function through Stochastic Gradient Descent

Denoting 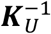 as ***S***_*U*_, and 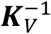 as ***S***_*V*_, the log-posterior of ***U*** and ***V*** is as follows:

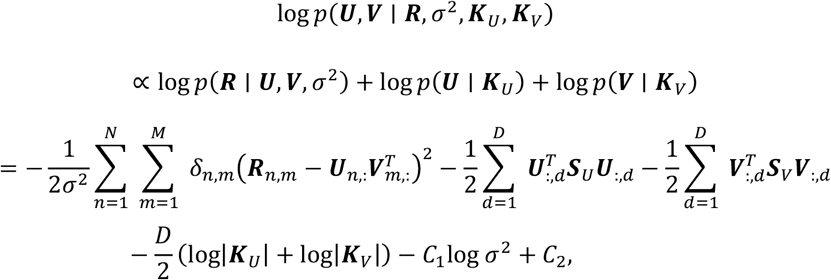

where *C*_1_ is the total number of non-missing entries in ***R***, and *C*_2_ is a constant independent of ***U*** and ***V***.

Therefore, maximizing the log-posterior above is equivalent to minimizing the following objective function:

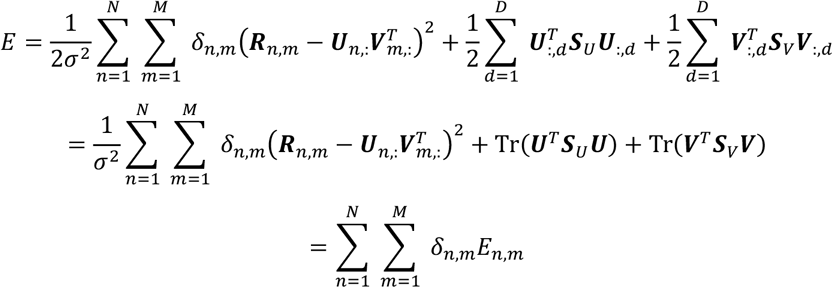

Then, we apply the Stochastic Gradient Descent (SGD) method to minimize the objective function. Denoting the number of non-missing entries in row *k* by *I*_*k*_, the gradients of objective function *E*_*n,m*_ concerning ***U***_*n*,:_ and ***V***_*m*,:_ are as follows:

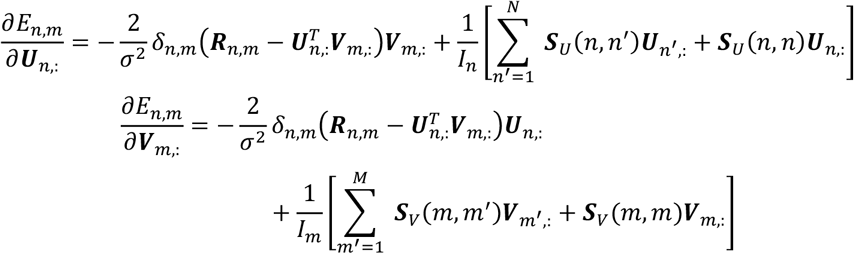

Finally, the updating rules for searching Maximum a Posteriori (MAP) estimation of ***U*** and ***V*** are as follows:

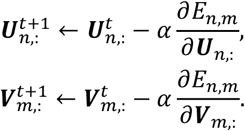

### Performance evaluation on multiple datasets

#### The prediction performance for non-missing entries

First, we divide the dataset into training and validation sets, employing stratified sampling based on the clonal information in either a 1:1 or 4:1 ratio. A smaller ratio is applied to the LT-scSeq datasets with relatively dense clonal information (CellTagging, *C. elegans*, and TraCe-seq), while a larger ratio is applied to those with relatively sparse clonal information (LARRY and Watermelon). Then we mask the entries in the validation set within the transition matrix ***R*** and use scTrace+ to approximately decompose the matrix ***R*** into matrices ***U*** and ***V***, so that the completed matrix ***R*\*** can be obtained through the product ***UV***^**T**^.

The performance of scTrace+ is evaluated from multiple perspectives (Fig. 3A).

Firstly, we directly compare the completed matrix ***R*\*** with the cell-cell clonal relationship matrix ***L***_*acr*_, and then calculate the AUC (Area Under the Receiver Operating Characteristic curve) scores. In detail, we posit that the raw lineage tracing data captures a portion of the real time-series relationships between cells across time points; thus, we consider entries equal to 1 in the matrix ***L***_*acr*_ as ground truth, with the expectation that the predicted fates would accurately identify these entries. The AUC scores are calculated based on the ROC curves, which plot the true positive rate (TPR) against the false positive rate (FPR) at different classification thresholds. Considering the high sparsity of ***L***_*acr*_, which can lead to class-imbalanced issues, we randomly split the positive samples in the validation set of ***L***_*acr*_ into five equal parts. For each part, we sample an equal number of negative samples, resulting in five sets of non-overlapping true values. Based on the sampled ground truth and the corresponding predicted values in ***R*\***, we compute the AUC score for five times.

As a data enhancement algorithm, we then select three performance indicators for scTrace+ when predicting non-missing entries:

1. Root Mean Square Error (RMSE): We calculate and record the RMSE for the validation set at each epoch, computed as 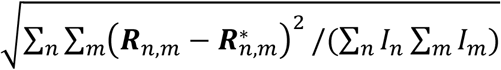;
2. Correlation: We conduct a correlation analysis between the predicted values and the ground truth of non-missing entries in the validation set. This analysis help us assess whether the model is overly biased towards predicting positive results;
3. Recall: We define a validation recall strictly. The threshold for extracting non-missing entries as positive samples is adaptive to the data distribution of training set. Specifically, if the true or predicted value of the validation set exceeds min(*μ*_*train*_ − 2 **\*** *σ*_*train*_, 0.5 **\*** max_*train*_), where 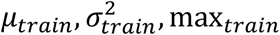 are the mean, variance, and maximum values of the non-missing entries in the training set, it is considered that a strong fate connection exists between the two cells in the corresponding row and column of the transition matrix. Based on this threshold, we record the recall for strong positive samples in validation sets during the training process.

The performance evaluation is tested on both developmental datasets (LARRY, CellTagging, and *C. elegans*) and tumor drug response datasets (Watermelon and TraCe-seq).

#### Ablation experiment for side information

To verify the effectiveness of integrating side information, we conduct ablation experiments on the datasets mentioned above. We compare three ablation scenarios (no side information, only the pre-timepoint side, and only the post-timepoint side) with the case of adding all side information. The evaluation is performed using the metrics of recall rates when predicting non-missing entries, as previously described.

#### Benchmarking against state-of-the-art methods

Here, we select the recent state-of-the-art method Cospar^7^ to benchmark the performance of scTrace+. Firstly, we generate the cell-cell transition matrix inferred by using both scTrace+ and Cospar. Then, we calculate AUC scores to evaluate and compare their performances (Fig. 4H and 5G).

Besides, we assess the accuracy rate of the inferred cell fates by scTrace+ and Cospar (Fig. 4H). For datasets that included cell type annotations from the original studies, we pre-define a set of reasonable cell-type level fates based on prior biological knowledge. We define the accuracy rates as the fraction of cells for which the predicted cell-type level fates (as detailed in the subsequent section, “Inferring cluster-level fate for each cell”) belonging to the pre-defined set of cell-type fates.

Additionally, we compare the proportions of cell-type level fate derived from the raw lineage information with those predicted by scTrace+ or Cospar (as detailed in the subsequent section, “Inferring cluster-level fate for each cell”). Then, we utilize cross-entropy to measure the divergence between the predicted fate proportion vector and those derived from the raw lineage information (Fig. S6A). This analysis allows us to determine which method, scTrace+ or Cospar, produces predictions that more closely reflect the randomness of fates indicated by the raw lineage tracing information (Fig. S6B and S8).

### Inferring cluster-level fate for each cell

To dissect the biological mechanisms underlying cell dynamics, we cluster the cells based on their expression profiles at each time point individually, and then assign the cluster-level fate to each cell based on the original lineage relationship matrix ***L***_*acr*_ and the enhanced transition matrix ***R*\***, respectively.

In the cell-cell lineage relationship matrix ***L***_*acr*_, each row can be seen as indicting the potential cell-level fates of each cell at pre-timepoint. To investigate the cell fate at the cluster-level, we merge “cell-to-cell” matrix ***L***_*acr*_ into a “cell-to-cluster” fate matrix ***F*** ∈ ℝ^*N,n*_*cluster*^ based on the clustering result at post-timepoint, so that each row in ***F*** denotes the fate vectors of the corresponding cells at pre-timepoint. Further, we define the dominant fate of a cell at pre-timepoint as the cluster at the post-timepoint that contains the highest number of cells sharing identical clonal information with that cell, corresponding to the argument of the maxima (abbreviated as argmax) in each row of ***F***.

For the enhanced transition matrix ***R*\***, we first filter out low values in ***R*\*** based on an adaptive threshold. In detail, we extract the entries in ***R*\*** whose corresponding pre-cell and post-cell share a common lineage barcode (belonging to the same clone), and set the threshold as the lower bound of these entries, excluding some outliers (less than *Q*_1_ − 1.5 **\*** (*Q*_3_ − *Q*_1_), where *Q*_1_ and *Q*_3_ are the first and third quartiles of the distribution). Then, we regard each row in ***R*\*** as the potential possibilities of a corresponding cell transitioning to other cells at post-timepoint, and further infer a cluster-level dominant fate for each cell at pre-timepoint (Fig. 4E). For each cell at pre- timepoint, we collect its transition values in ***R*\*** to a selected post-cluster *k* and to all other post-clusters *k*^−^, respectively. Then, we perform a one-sided Wilcoxon rank-sum test or Mann–Whitney test (using the ranksums or mannwhitneyu functions in the Scipy package) for a ‘greater than’ comparison between these two groups of transition values. The alternative hypothesis is that transition values to cluster *k* are more likely to be larger than those transitioning to *k*^−^. For all post-clusters, we obtain a list of p-values and assign the cluster with the minimum p-value as the dominant fate for the current cell, when the sufficient statistical significance is satisfied min{*p*_*k*_, *k* **=** 1, 2, …, *n*_*post*_*clusters*_} < 0.05. Finally, we integrate the original lineage barcoding information with the inferred dominate fates as the enhanced cluster-level cell fates.

### Intra-cluster dynamic heterogeneities analysis

To explore the intra-cluster dynamic heterogeneities, we perform differential expression analysis between the subclusters with distinct fates based on both the raw and the enhanced cluster-level cell fates. Differentially expressed genes (DE genes) are identified by the Wilcoxon rank-sum test in the Scanpy package. Then, we compare the DE genes detected before and after data enhancement to evaluate whether scTrace+ facilitates some novel findings about cell fate decision (Fig. 4F, 4G and 5D).

In the TraCe-seq dataset, we further analyze the effect of these DE genes on patient survival. We collect the bulk gene expression and patient survival data from TCGA-LUAD (lung adenocarcinoma) cohorts, which is consistent with the tumor type of TraCe-seq data. Then, we fit a Cox proportional hazards regression model to examine the relationship between the survival information and the expression levels of each DE gene. A hazard ratio calculated greater than 1 indicates an increased hazard associated with this DE gene, while a value less than 1 suggests a decreased hazard (Fig. 5E). Besides, the DE genes are further used to perform enrichment analysis on the cancer hallmark pathways in MSigDB^31^ to identify the novel signaling pathways facilitated by the fate enhancement (Fig. 5F).

Notably, for datasets that provide the cell type labels in the original study, we use the cell types instead of cell clusters for the above analyses.

