## Supplementary Figures for "scTrace+: enhance the cell fate inference by integrating the lineage-tracing and multi-faceted transcriptomic similarity information"

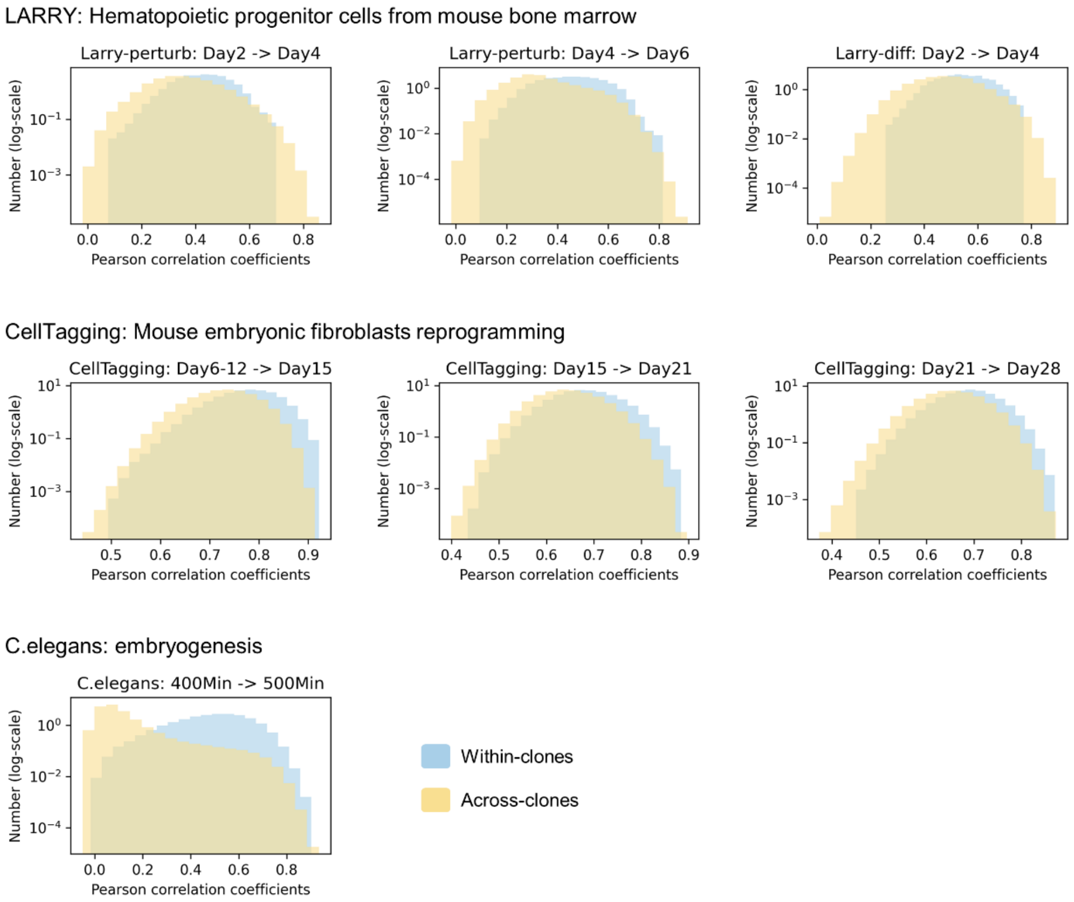


**Fig. S1. Comparison between transcriptomic similarity and lineage information across time points in developmental datasets (related to Fig. 1D).**

Histograms comparing the transcriptomic similarity among cells within the same clone (blue) versus those from different clones (yellow) across time points in developmental datasets, including: LARRY-diff, LARRY-perturb, CellTagging, and C.elegans.

* In all the subfigures of **Fig.S1~Fig.S4**: The x-axis “Pearson correlation coefficients” refers to “transcriptomic similarity” in **Fig.1D**, which is generated by “corrcoef” in the numpy package. The y-axis “Number (log-scale)” refers to “Density (log-scale)” in **Fig.1D**.

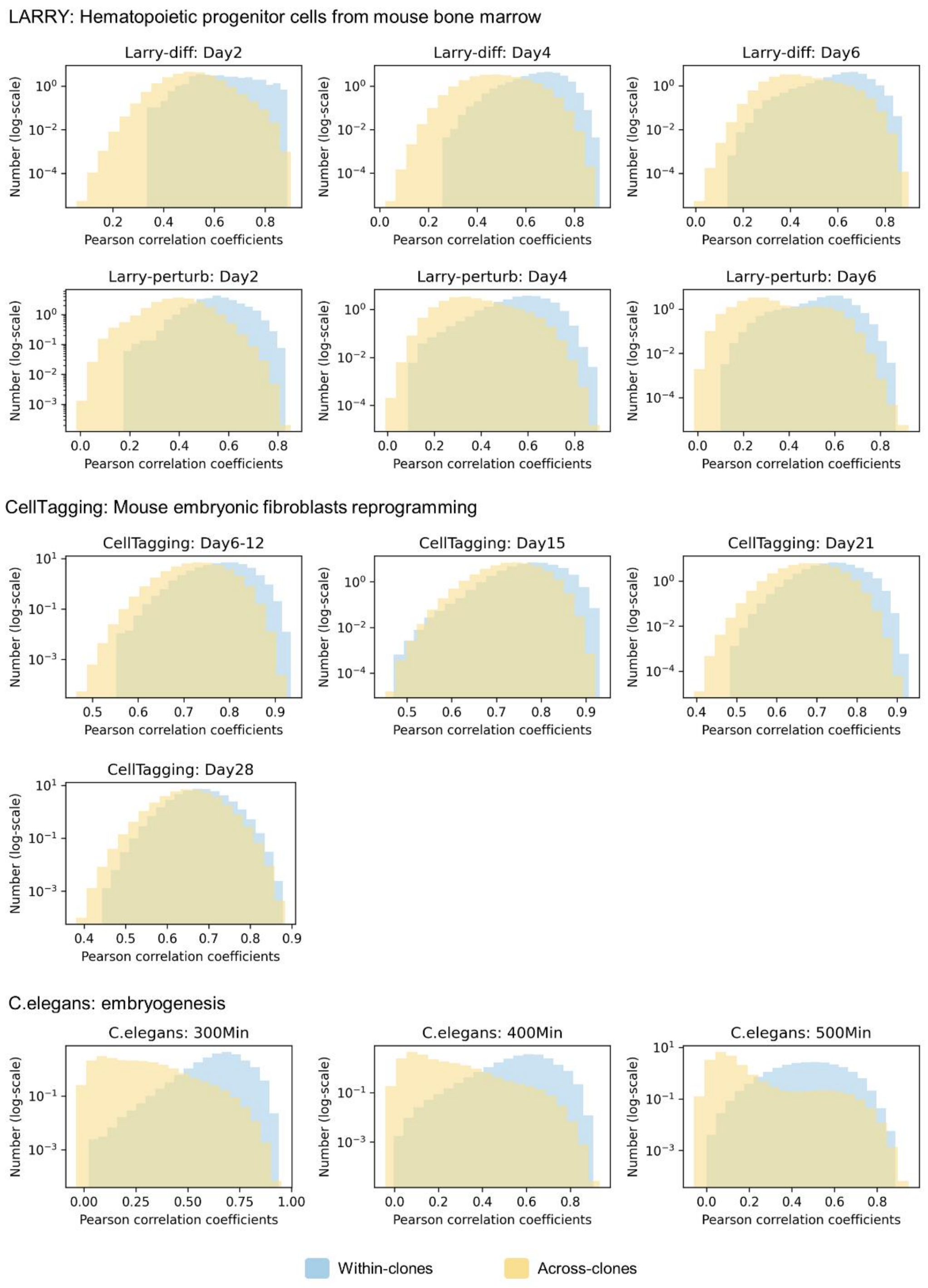


**Fig. S2. Comparison between transcriptomic similarity and lineage information within a single time point in developmental datasets (related to Fig. 1D).**

Histograms comparing the transcriptomic similarity among cells within the same clone (blue) versus those across different clones (yellow) at each time point in developmental datasets, including: LARRY-diff, LARRY-perturb, CellTagging, and C.elegans.

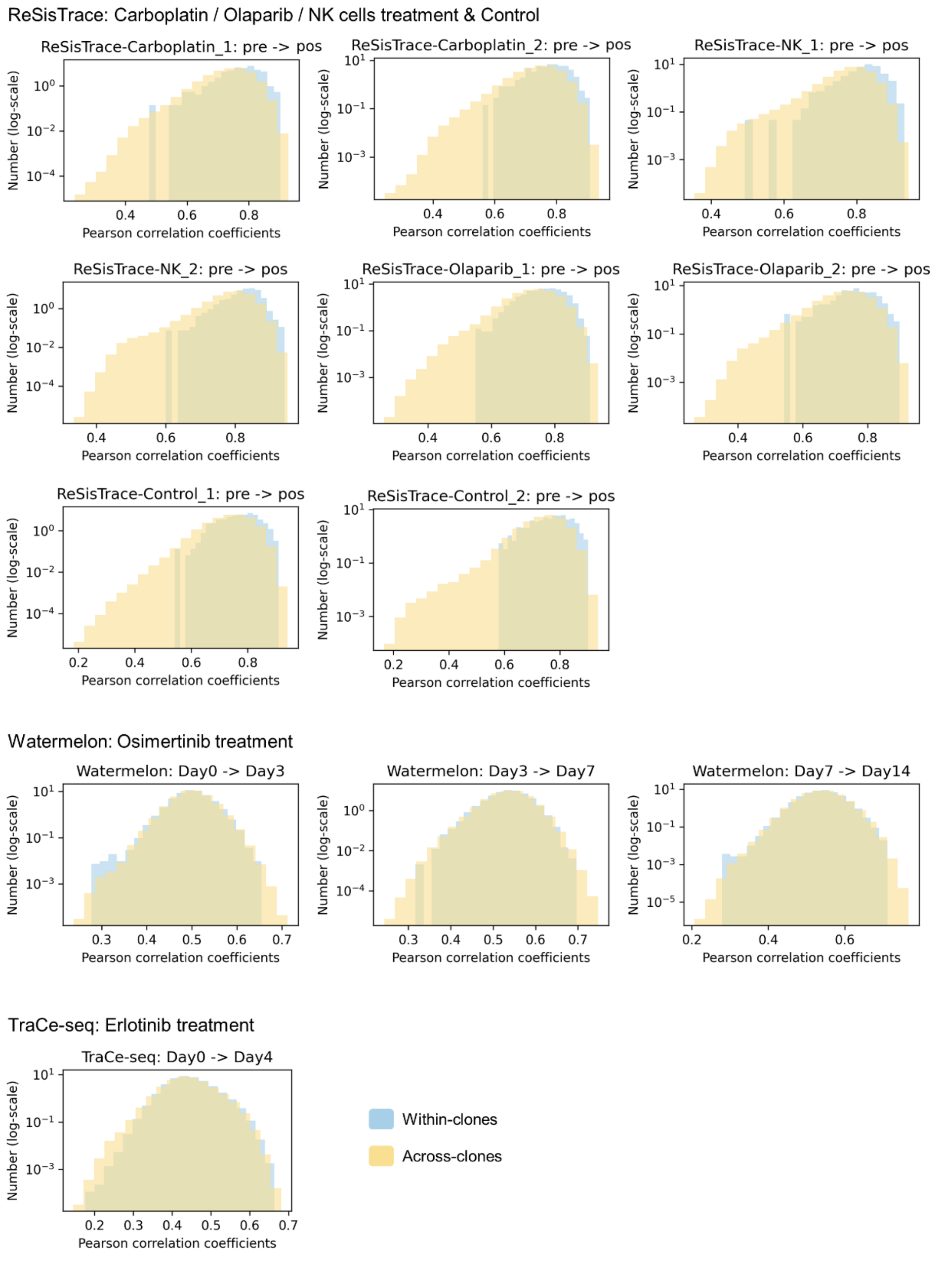


**Fig. S3. Comparison between transcriptomic similarity and lineage information across time points in drug-perturbed tumor datasets (related to Fig. 1D).**

Histograms comparing the transcriptomic similarity among cells within the same clone (blue) versus those from different clones (yellow) across time points in drug-perturbed tumor datasets, including: ReSisTrace, Watermelon, and TraCe-seq.

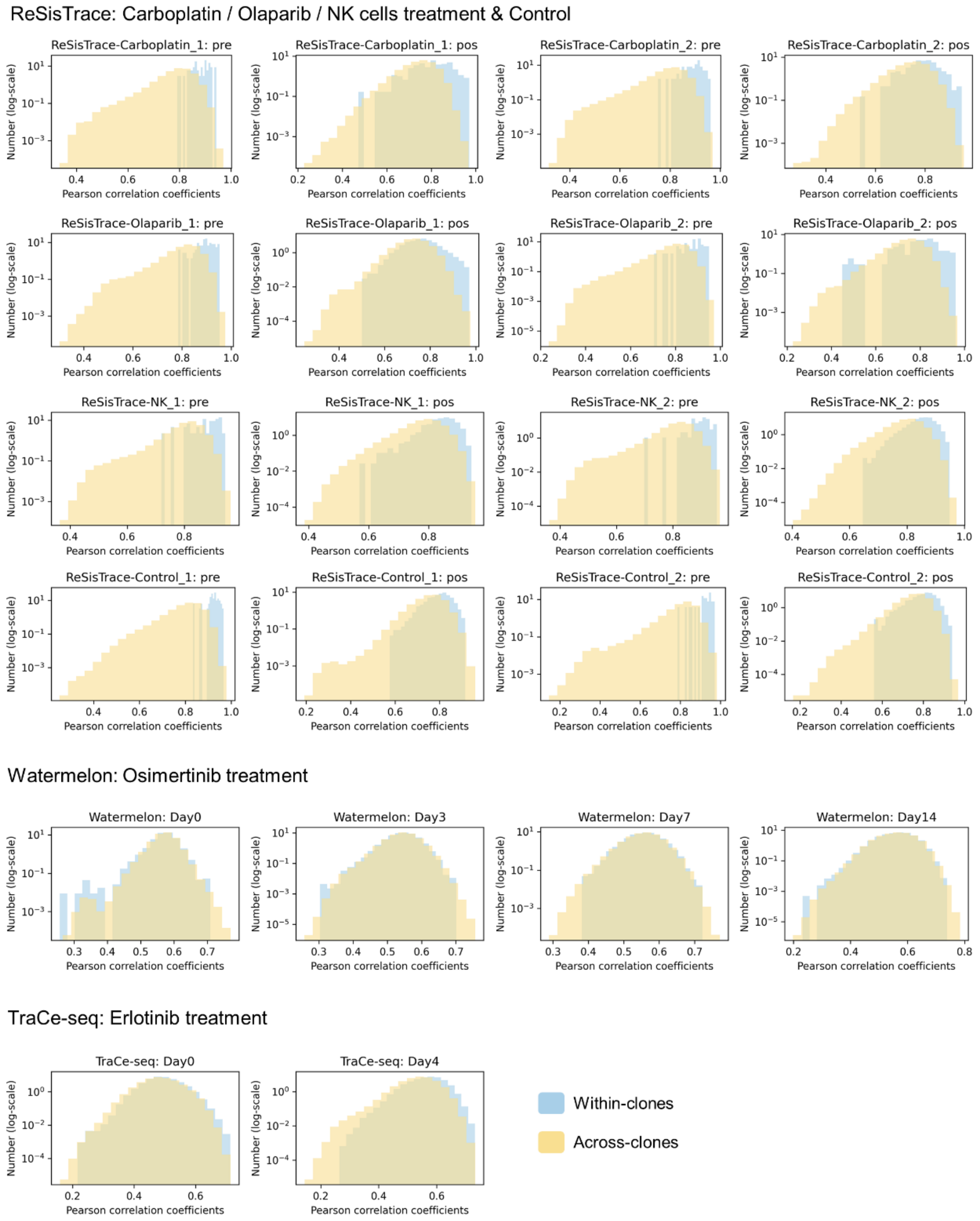


**Fig. S4. Comparison between transcriptomic similarity and lineage information within a single time point in drug-perturbed tumor datasets (related to Fig. 1D).** Histograms comparing the transcriptomic similarity among cells within the same clone (blue) versus those across different clones (yellow) at each time point in drug-perturbed tumor datasets, including: ReSisTrace, Watermelon, and TraCe-seq.

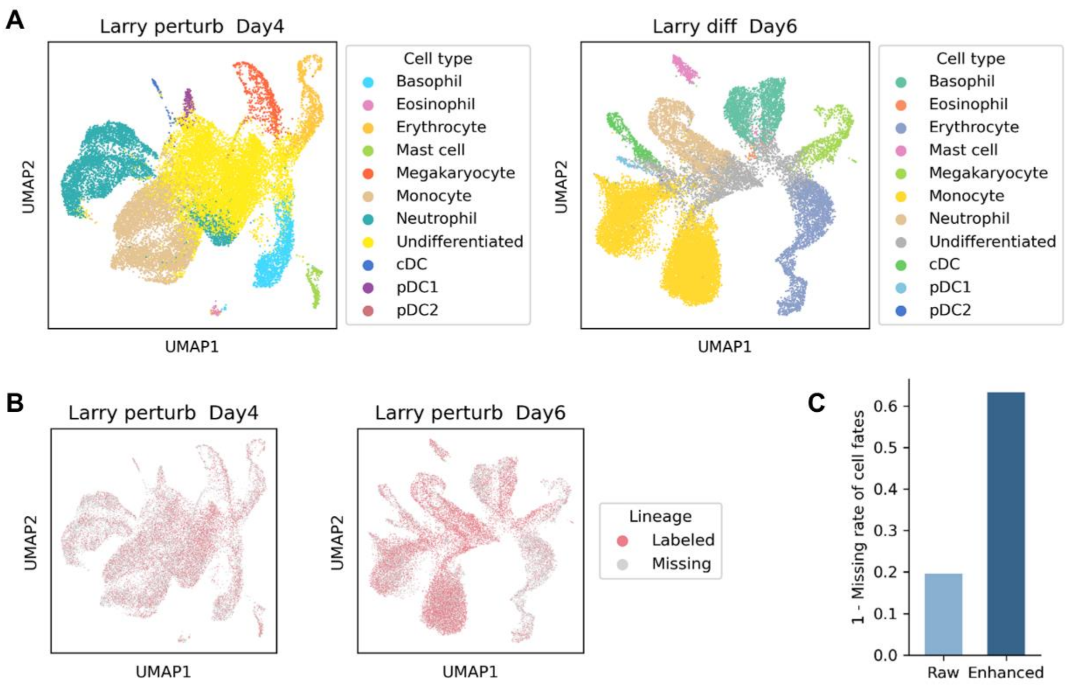


**Fig. S5. The cell type annotation results and the missing profile of cell fates in Larry-perturb dataset (related to Fig. 4).**

**A.** UMAP plots of the cells in Larry-perturb dataset at days 4 and 6. The color indicates the cell type label annotated in the original study. **B.** UMAP plots showing the lineage barcode labeling results for the cells in Larry-perturb dataset. **C.** The missing rates of cell fates before (light blue) and after (dark blue) the enhancement of scTrace+ in Larry-perturb dataset.

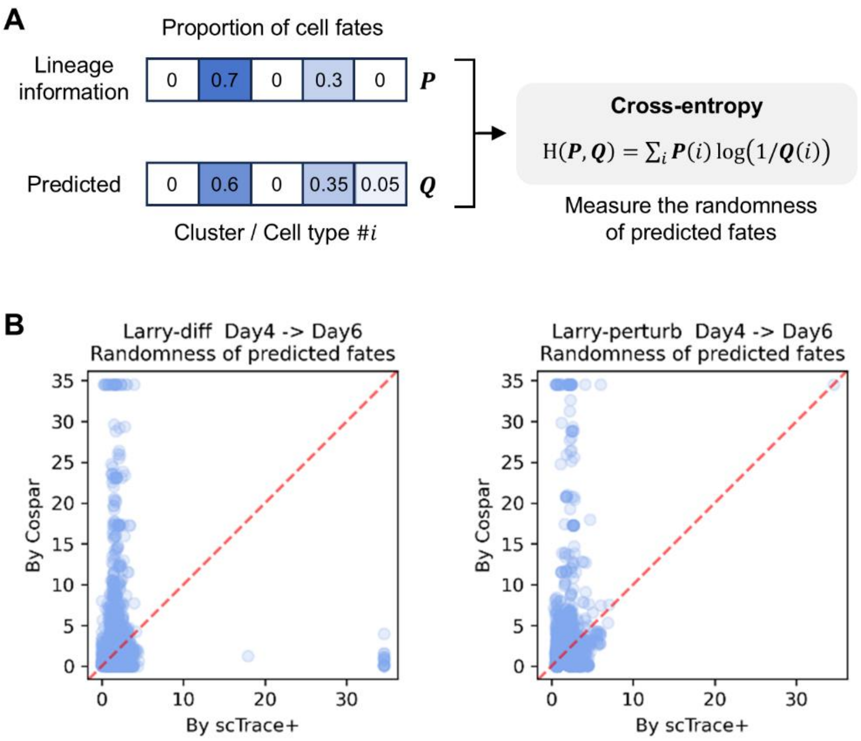


**Fig. S6. The randomness of the predicted fates in Larry-diff and Larry-perturb datasets (related to Fig. 4).**

**A.** Schematic diagram showing the approach of measuring the randomness of the predicted fates, which is defined as the cross-entropy between the predicted cluster-level (or cell type level) fate proportion vector by enhancement algorithm (scTrace+ or Cospar) and those derived from the raw lineage information. **B.** Scatter plots to compare the randomness of cell fates predicted by scTrace+ (x-axis) and Cospar (y-axis) in Larry-diff (left panel) and Larry-perturb (right panel) datasets. The red dashed lines indicate equal randomness.

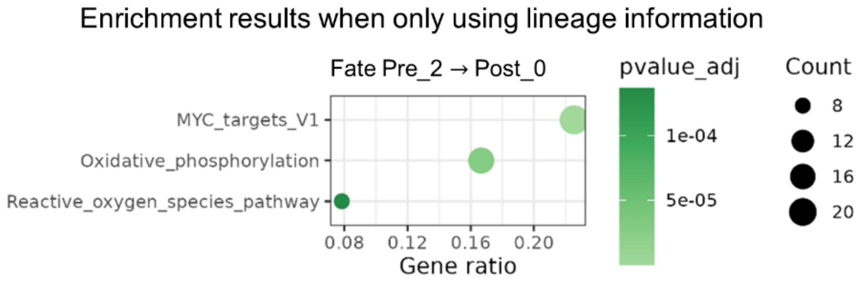


**Fig. S7. The pathway enrichment analysis results for the DE genes of fate Pre_2→ Post_0 before enhancement in TraCe-seq dataset (related to Fig. 5F).** The colors of points indicate the adjusted $p\_value$, and the size of points reflects the number of genes belonging to the corresponding pathway.

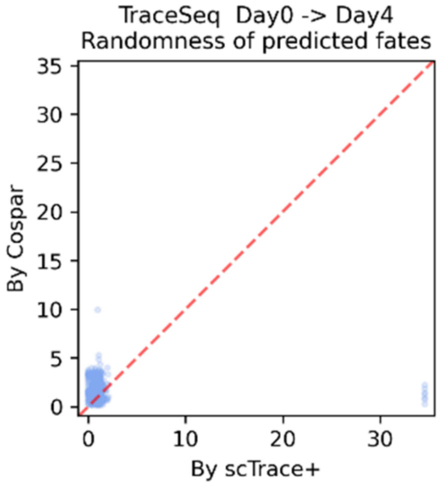


**Fig. S8. Comparison between the randomness of cell fates predicted by scTrace+ and Cospar (related to Fig. 5).** Scatter plots to compare the randomness of cell fates predicted by scTrace+ (x-axis) and Cospar (y-axis) in TraCe-seq dataset. The red dashed lines indicate equal randomness.
