## Supplementary Tables for "scTrace+: enhance the cell fate inference by integrating the lineage-tracing and multi-faceted transcriptomic similarity information"

**Supplementary Table 1:** Algorithm comparison.

| **Algorithm** | **Lineage relationships** | | **Transcriptomic similarities** | |
|---|---|---|---|---|
| | **Within-timepoint** | **Across-timepoints** | **Within-timepoint** | **Across-timepoints** |
| **LineageOT** | √ | | | √ |
| **Moslin** | √ | | | √ |
| **Cospar** | √ | | √ | |
| **scTrace+** | √ | √ | √ | √ |

**Supplementary Table 2:** Dataset information.

| Dataset | Time Points | Source | Category & Description |
|---|---|---|---|
| Larry-diff[1] | Day2, Day4, Day6 | https://github.com/AllonKleinLab/paper-data/ | Hematopoietic progenitor cells from mouse bone marrow, In vitro differentiation time course |
| Larry-perturb[1] | Day2, Day4, Day6 | https://github.com/AllonKleinLab/paper-data/ | Hematopoietic progenitor cells from mouse bone marrow, In vitro cytokine perturbations |
| C.elegans[2] | 300 minutes, 400 minutes, 500 minutes | GSE126954 | *C. elegans* embryogenesis |
| CellTagging[3] | Day (6, 9, 12) *, Day 15, Day 21, Day28 | GSE99915 | Mouse embryonic fibroblasts reprogramming |
| Watermelon[4] | Day 0, Day 3, Day 7, Day 14 | https://github.com/yaaraore/watermelon | Osimertinib-perturbed cancer cell line |
| TraCe-seq[5] | Day 0, Day 4 | E-MTAB-10698 | Erlotinib-perturbed PC9 cancer cell line |
| ReSisTrace[6] | Pre-treatment, Pos-treatment | GSE223003 | Perturbed (Carboplatin, Olaparib, NK cells) cancer cell line |

* Day 6-12 were integrated according to the original paper (TF delivery before reprogramming): "Tagging with a second library (CellTagD3) was performed at the end of the 3-day period of transcription factor delivery, followed by a third round (CellTagD13) 13 days after the start of reprogramming"
